# A comparison of auditory oddball responses in dorsolateral prefrontal cortex, basolateral amygdala and auditory cortex of macaque

**DOI:** 10.1101/326280

**Authors:** Corrie R. Camalier, Kaylee C. Scarim, Mortimer Mishkin, Bruno B. Averbeck

**Author notes:** Correspondence: Corrie R. Camalier, 49 Convent Drive, MSC 4415, Bethesda, MD 20892-4415.

## Abstract

The mismatch negativity (MMN) is an event-related potential component seen in response to unexpected “novel” stimuli, such as in an auditory oddball task. The MMN is of wide interest and application, but the neural responses that generate it are poorly understood. This is in part due to differences in design and focus between animal and human oddball paradigms. For example, one of the main explanatory models, the “predictive error hypothesis”, posits differences in timing and selectivity between signals carried in auditory and prefrontal cortex. However these predictions have not been fully tested because 1) noninvasive techniques used in humans lack the combined spatial and temporal precision necessary for these comparisons, and 2) single-neuron studies in animal models, which combine spatial and temporal precision, have not focused on higher order contributions to novelty signals. In addition, accounts of the MMN traditionally do not address contributions from subcortical areas known to be involved in novelty detection, such as the amygdala. To better constrain hypotheses and to address methodological gaps between human and animal studies, we recorded single neuron activity from the auditory cortex, dorsolateral prefrontal cortex, and basolateral amygdala of two macaque monkeys during an auditory oddball paradigm modeled after that used in humans. Consistent with predictions of the predictive error hypothesis, novelty signals in prefrontal cortex were generally later than in auditory cortex, and were abstracted from stimulus-specific effects seen in auditory cortex. However, we found signals in amygdala that were comparable in magnitude and timing to those in prefrontal cortex, and both prefrontal and amygdala signals were generally much weaker than those in auditory cortex. These observations place useful quantitative constraints on putative generators of the auditory oddball-based MMN, and additionally indicate that there are subcortical areas, such as the amygdala, that may be involved in novelty detection in an auditory oddball paradigm.

## Introduction

An organism’s ability to extract patterns from the world, and to quickly detect deviation from these predictions, is key to survival (Friston, 2009). The auditory oddball task is one of the dominant paradigms used to examine mechanisms of novelty detection (Naatanen et al., 2012; Nelken, 2014). In this task, an auditory stimulus is presented repeatedly as a “standard” and is infrequently interleaved with a different “deviant” or “oddball” auditory stimulus. The response to the deviant is greater than the response to the standard, or to the response of the deviant stimulus when played as a standard. This early difference in response is referred to as the “mismatch negativity” (MMN) in scalp-based event related potentials. Though the neural basis of this potential is poorly understood, the MMN is of wide use and interest as an index of early sensory processing and novelty sensitivity, and is of clinical interest as a potential index of symptoms of some psychiatric disorders.

To examine the basis of the auditory oddball mediated MMN, human studies using noninvasive measures (fMRI, EEG, MEG) have suggested that it has both temporal and frontal cortical generators. According to the predictive error hypothesis of the MMN, sensory and frontal generators have distinct functions (Garrido, Kilner, Stephan, & Friston, 2009; Naatanen, Teder, Alho, & Lavikainen, 1992). While the standard is being repeated, a stimulus specific memory is built up in sensory (auditory) cortex, and this memory drives an expectation. On the other hand, frontal cortex represents whether there is a difference between the expected and actual stimulus (Garrido et al., 2009; Giard, Perrin, Pernier, & Bouchet, 1990; Naatanen et al., 1992). This hypothesis predicts that frontal novelty signals should arise later than those in auditory cortex, and additionally that the signal in prefrontal cortex be abstracted from any stimulus selective activity seen in auditory cortex. It has been difficult to test this hypothesis in humans using noninvasive techniques because they do not combine the precise temporal and spatial resolution needed to compare timecourses with certainty (Deouell, 2007; Rinne, Alho, Ilmoniemi, Virtanen, & Naatanen, 2000; Tse & Penney, 2008). In addition, to compare stimulus specificity, one must be able to compare the novelty response between stimulus types across areas, which is not commonly done. Human electrocortocogtraphy (ECoG) studies have also been used to examine the correlates of oddball and novelty detection, and have confirmed frontal and sensory components. However, heterogeneity of clinical electrode placement within and between patients makes it difficult to systematically compare magnitude and timecourse of early novelty signals between areas (Durschmid, Edwards, et al., 2016; Edwards, Soltani, Deouell, Berger, & Knight, 2005; Rosburg et al., 2005). In contrast, animal models of auditory oddball paradigms do have the necessary spatial and temporal specificity to compare signals across areas. However, these studies have primarily been concerned with mechanisms of oddball in early auditory processing (cochlear nucleus through auditory cortex), and not activity from other areas (Ayala, Perez-Gonzalez, Duque, Nelken, & Malmierca, 2012; Fishman, 2014; Javitt, Steinschneider, Schroeder, Vaughan, & Arezzo, 1994; Parras et al., 2017; Ulanovsky, Las, & Nelken, 2003; Yarden & Nelken, 2017). In addition, the majority of animal studies choose neuron-specific “standard” and “deviant” stimuli based on the receptive fields of the neuron being recorded at the time. This tuned design has been undeniably powerful for other questions (e.g. mechanisms of adaptation and effects of pharmacological manipulations at a single-neuron level), but may not be ideal for connecting to the larger body of human literature where, by definition, identical standard and deviant stimuli are used for all areas. This distinction appears especially important when questions pertain to the comparison of timecourse and magnitude of novelty signal between areas.

Largely separate from accounts involving the cortical substrates of the MMN in oddball, a body of work has established correlates of novelty and salience detection in subcortical nuclei such as the amygdala and nucleus accumbens (NAc) (Balderston, Schultz, & Helmstetter, 2013; Blackford, Buckholtz, Avery, & Zald, 2010; Bradley et al., 2015; Zaehle et al., 2013). For example, a human intracranial study suggested robust auditory oddball-based MMN in NAc (Durschmid, Zaehle, et al., 2016). Amygdala activation in auditory oddball has been reported in at least one fMRI study (Czisch et al., 2009), but an intracranial study suggested that auditory oddball novelty signal may only arise in the amygdala during active detection (Kropotov et al., 2000). There has thus been renewed interest in these as nuclei that can affect either directly or indirectly, the MMN, with the understanding that source localization techniques used in human noninvasive studies can miss subcortical generators. However, it is unclear whether these areas’ signals are as fast or as robust as those of the hypothesized cortical generators. The human intracranial studies described above are suggestive, but cannot systematically compare the signals in amygdala to that of other areas due to the necessary heterogeneity of clinical placement. Studies in animal models would be able to address magnitude and timing differences between these and more established generators but extant studies have largely focused on examining oddball responses in early auditory areas. The goal of the present experiment, therefore, was to systematically compare the single neuron correlates of an early auditory oddball signal in auditory cortex, dorsolateral prefrontal cortex and basolateral amygdala. This approach, drawing on a substantial number of neurons (~600 per area afforded by a multichannel approach), allows us to address predictions of the predictive error account in auditory and prefrontal cortex and evaluate whether the account should be extended to areas such as the amygdala. This comparison, with the advantage that it is modeled after human studies, provides useful quantitative constraints for future accounts of the brain bases of auditory oddball-mediated MMN.

## Materials and Methods

### Subjects, surgical procedures, and neurophysiological data acquisition

Experiments were carried out on two adult male rhesus macaques (*Macaca mulatta*). All experimental procedures were approved by the National Institute of Mental Health Animal Care and Use Committee and followed *Guide for the Care and Use of Laboratory Animals.* Before data acquisition, monkeys were implanted with titanium head posts for head restraint. In a separate procedure, monkeys were fitted with custom acrylic chambers oriented to allow vertical grid access to the left dorsolateral prefrontal cortex (dorsal bank of the principal sulcus extending ventral, >1 mm away from arcuate sulcus, roughly 46/8Ad), the lateral portion of the amygdala (entire dorsoventral extent, primarily basolateral amygdala), and auditory cortex (primarily A1 but including small portions of lateral belt areas). Recording areas were verified though a T1 scan of grid coverage with respect to underlying anatomical landmarks (Fig. 1B-C), combined with maps of frequency reversals and response latencies of single neurons to determine A1 location and extent (Camalier, D’Angelo, Sterbing-D’Angelo, de la Mothe, & Hackett, 2012). Recordings were made using either 16 or 24 channel laminar “V-trodes” (Plexon, Inc, Dallas TX; 200-300 μm contact spacing, respectively), which allowed identification of white matter tracts, further allowing identification of electrode location with respect to sulci and gyri. Electrodes were advanced to their target location (NAN microdrives, Nazareth, Israel) and allowed to settle for at least 1 hour before recording.

Multichannel spike and local field potential recordings were acquired with a 64 channel data acquisition system. Spike signals were amplified, filtered (0.3-8 kHz), and digitized at ~24.4 kHz. Spikes were initially sorted online on all channels using real-time window discrimination. Digitized waveforms (snippets) and timestamps of stimulus events were saved for final sorting (Plexon offline sorter V 3.3.5). The units were also graded according to isolation quality (single or multi neurons). Single and multiunits were analyzed separately, but the patterns of results were similar, and so were combined. The acquisition software interfaced directly with the stimulus delivery system and both systems were controlled by custom software (OpenWorkbench and OpenDeveloper, controlling a RZ2, RX8, Tucker Davis Technologies (TDT) System 3, Alachua, FL).

### Stimuli, experimental design, and statistical analysis

We utilized a “flip-flop” auditory oddball design used in intracranial studies (e.g. Naatanen, Gaillard, & Mantysalo, 1978; Ulanovsky et al., 2003). This paradigm controls for differences in activity driven by the stimulus selectivity of neurons unrelated to stimulus type, where type is standard or deviant (Fig. 1A). Since our questions were primarily concerned with comparisons of neural responses in auditory cortex and higher-order areas, the stimuli used were constant across all areas and neurons and chosen to drive responses in higher order areas such as the amygdala and prefrontal cortex, and thus were more complex than pure tones. The stimuli, 300 ms 1 kHz and 8 kHz square waves, are perceptually distinct, spectrally nonoverlapping wideband stimuli likely to activate large parts of the auditory cortex. The spectrum of a square waves contains the odd harmonics of the fundamental frequency, such that the 1kHz square wave contains 1kHz and harmonics of 3, 5, 7, 9, up to ~33 kHz, and the 8 kHz square wave contains 8 kHz and a functional harmonic of 24 kHz, as macaques can hear up to about 35 kHz (Hauser, Burton, Mercer, & Ramachandran, 2018; Jackson, Heffner, & Heffner, 1999; Recanzone, Guard, & Phan, 2000). These stimuli have minimal spectral overlap, which is thought to be important for evoking MMN correlates (see Khouri & Nelken, 2015). The sounds were 300 ms in duration (with 5 ms cos^2^ onset/offset ramps to minimize spectral splatter) and they were presented from a speaker 10 cm from the contralateral (right) ear calibrated to 60 dB SPL.

The “flip-flop” design has two blocks, one in which the 1 kHz pulse is the standard and the 8 kHz the deviant, and the other in which standard and deviant identities are reversed. The flip-flop design critically allows the ability to dissociate any stimulus specificity of the response from actual novelty; the measure of most interest is the difference between the response to a stimulus when it is presented as a standard vs when it is a presented as a deviant (DEV-STD). If this comparison is positive, then the response to a stimulus presented as a deviant is greater than when it is presented as a standard. Each block in the flip-flop design contained 270 standards and 30 deviants (10% deviant probability) presented in randomized order with an inter-onset interval between 760-860 ms. Ten additional standards were added to the beginning of each block to ensure a stable standard trace (Fig. 1). The deviant stimulus identity of the first block (1 kHz or 8 kHz) was switched across days to eliminate any order effects. During the oddball task, head-restrained monkeys sat quietly in a primate chair, watching a soundless video (common in human oddball studies) in a double-walled acoustically isolated sound booth (Industrial Acoustics Company, Bronx, NY).

**Figure 1.**
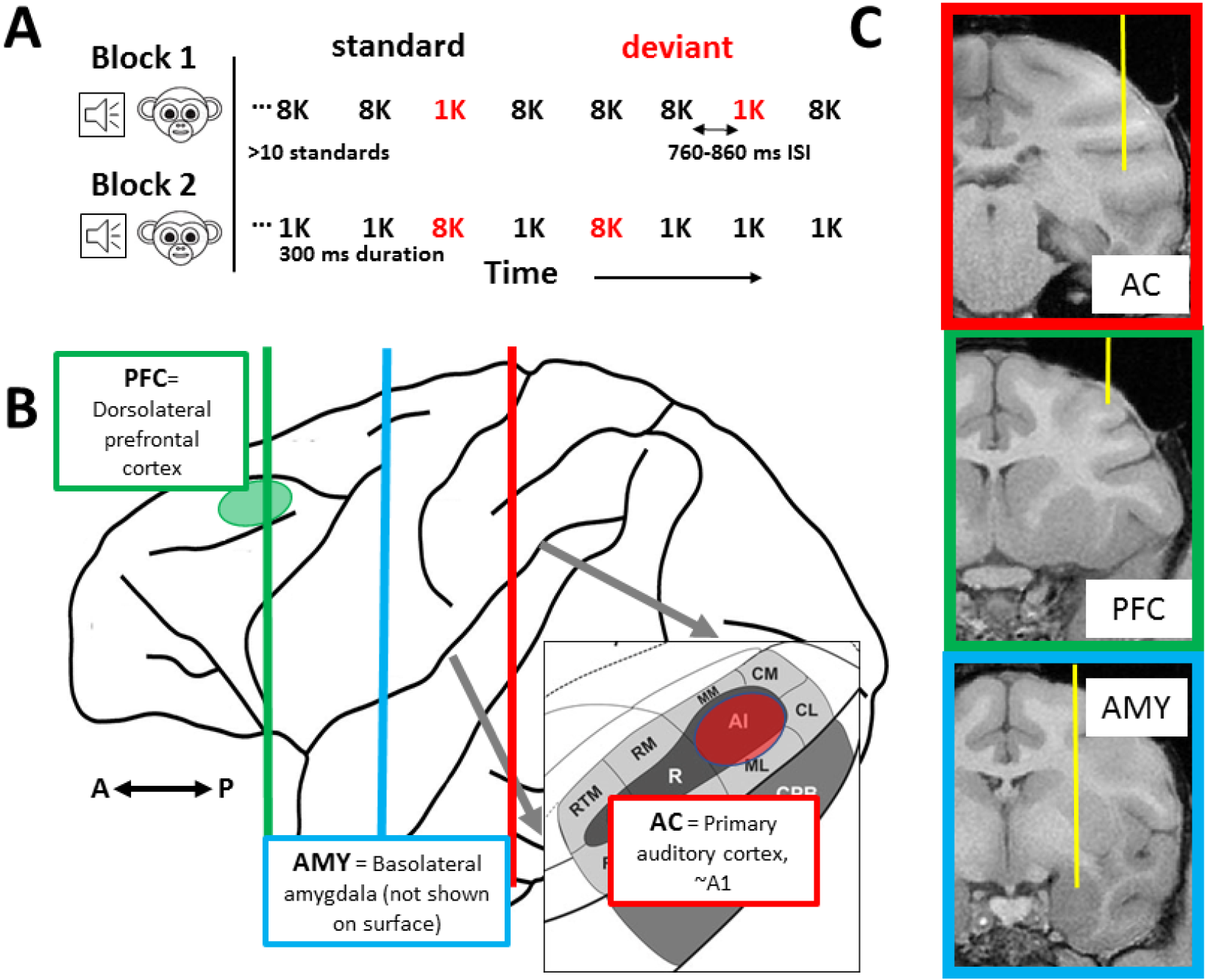
Task design, recording locations and trajectory. A. Details of the “flip-flop” oddball task. The 1 kHz and 8 kHz stimuli were consistently used for all areas. B. Schematic of macaque brain with locations of recording areas shown on surface (where possible) in red (AC) and green (dlPFC). Vertical lines indicate sagittal slices shown in panel C. C. Exemplar recording trajectory for each area reconstructed from T1-weighted scan - all areas were recorded with multichannel laminar probes oriented vertically which allowed for a comparison of ~600 neurons per area.

All data analysis was performed with custom scripts in Matlab (Natick, MA). Code and data are available upon reasonable request to the corresponding author. Data from the first 10 standards was discarded, and then spikes evoked by each stimulus (1 kHz vs 8 kHz) and type (standard vs deviant) combination were calculated over time. For all analyses, mean evoked spike rates were converted to z-scores using the baseline mean and standard deviation (-150-0 ms prestimulus).

### ANOVA models

To establish the population of auditory-responsive cells (here defined as responding to any of the stimuli in any context), we performed a 100 ms sliding window ANOVA on the evoked activity of each neuron (timebin advanced from 0-250 ms in 20 ms increments), including the factors of stimulus frequency (1 vs 8 kHz) and type (deviant vs standard) and their interaction. If any factor was significant in any window (p<0.01, FDR corrected for multiple comparisons), the neuron was considered task responsive. For the population-based analysis of oddball selectivity by area and stimulus (Fig. 4), we used an analysis of a mixed within-between effects ANOVA (3×2×2; area × stimulus × type). In this case, neuron identity was specified as a “random” effect in the model and was nested under area, where area is the region recorded from. To establish oddball selectivity at the level of individual neurons, if a neuron exhibited a significant response to type, and response to deviant > standard, or a significant interaction and one of the stimuli had a deviant response > standard response (p<0.01, Bonferroni corrected as appropriate), then it was considered to exhibit deviance selectivity analogous to the mismatch negativity. To be conservative, we refer to this firing-rate based deviance selectivity as simply “oddball selectivity”. It has the same pattern as the MMN measured by continuous methods such as EEG but is not derived from a continuous measure. Note this measure is similar to “stimulus-specific adaptation” (SSA) used in other reports, but as the basis of this is probably not strictly adaptation, especially in higher-order areas examined here, we prefer to use descriptive nomenclature to avoid confusion.

For the individual-neuron based analysis of time course of deviance detection (Fig. 6A), for each responsive neuron in each area, we measured the first timebin that exhibited a selective response to stimulus type (type x stim, 30 ms windows sliding at 10 ms (p<0.05, Bonferroni corrected)). For the population-based analysis of the time course of novelty detection (Fig. 6B-D), we used a sliding ANOVA (mixed within-between, stimulus x type x neuron identity) on response in 30 ms bins with 10 ms intervals and looked for periods with a significant effect of type (p<0.01, Bonferroni corrected). For the analysis of whether adaptation exists and is different between areas (Fig. 7), we calculated the evoked response as a function of time (either presentation number since the beginning of each block or presentation number following deviant) for each responsive neuron. To quantify this time course, we fit an exponential decay function (R=a*exp(b*t), where presentation #= t) (After Antunes, Nelken, Covey, & Malmierca, 2010; Ulanovsky, Las, Farkas, & Nelken, 2004) to each neuron’s time course for each stimulus, and then determined whether the population of decay constants (b) were less than 0 via a one-tailed t-test. If adaptation was indicated, ANOVAs on the population of b values was done to determine whether adaptation was greater for some stimuli. For the analysis of within block adaptation, this was done for the first 27 presentations. This covers the majority of the adaptation, and includes the same number of stimuli for standards as deviants. For the analyses relative to the last deviant, adaptation was calculated for the first 6 presentations of standards following a deviant. Consistent with a population-based approach done by other studies (see above), these results were confirmed by a bootstrap test on 1000 fits done to averages of 70 neurons (chosen randomly from population), and the results were identical.

## Results

We recorded neurons from primary auditory cortex (AC, n=690), dorsolateral prefrontal cortex (PFC, n=598) and the basolateral amygdala (AMY, n=627), while monkeys were presented with a flip-flop auditory oddball paradigm (Fig. 1A). In all three areas, we found neurons that responded more strongly to the stimulus when presented as an oddball (examples in Fig. 2). Note that generally both baseline firing rates and responses were weaker in amygdala and prefrontal cortex, so subsequent analyses focus on the normalized differences in firing rates expressed as a z-score. To compare novelty responsiveness and stimulus selectivity across areas, we first identified the fraction of neurons that were generally auditory responsive under these oddball task conditions. In auditory cortex 462 neurons were responsive to the task stimuli in any context (67%). Smaller proportions were responsive in prefrontal cortex (105, 18%) and amygdala (105, 17%). This analysis was done using a fixed window (0-250 ms) chosen to correspond to the period of MMN generation in the primate (Gil-da-Costa, Stoner, Fung, & Albright,2013). To ensure that this was not due to the choice of analysis window, we examined larger or smaller sliding windows (50, 200 ms) and the pattern of results did not change, though the fixed window yielded more conservative estimates of responsiveness.

**Figure 2.**
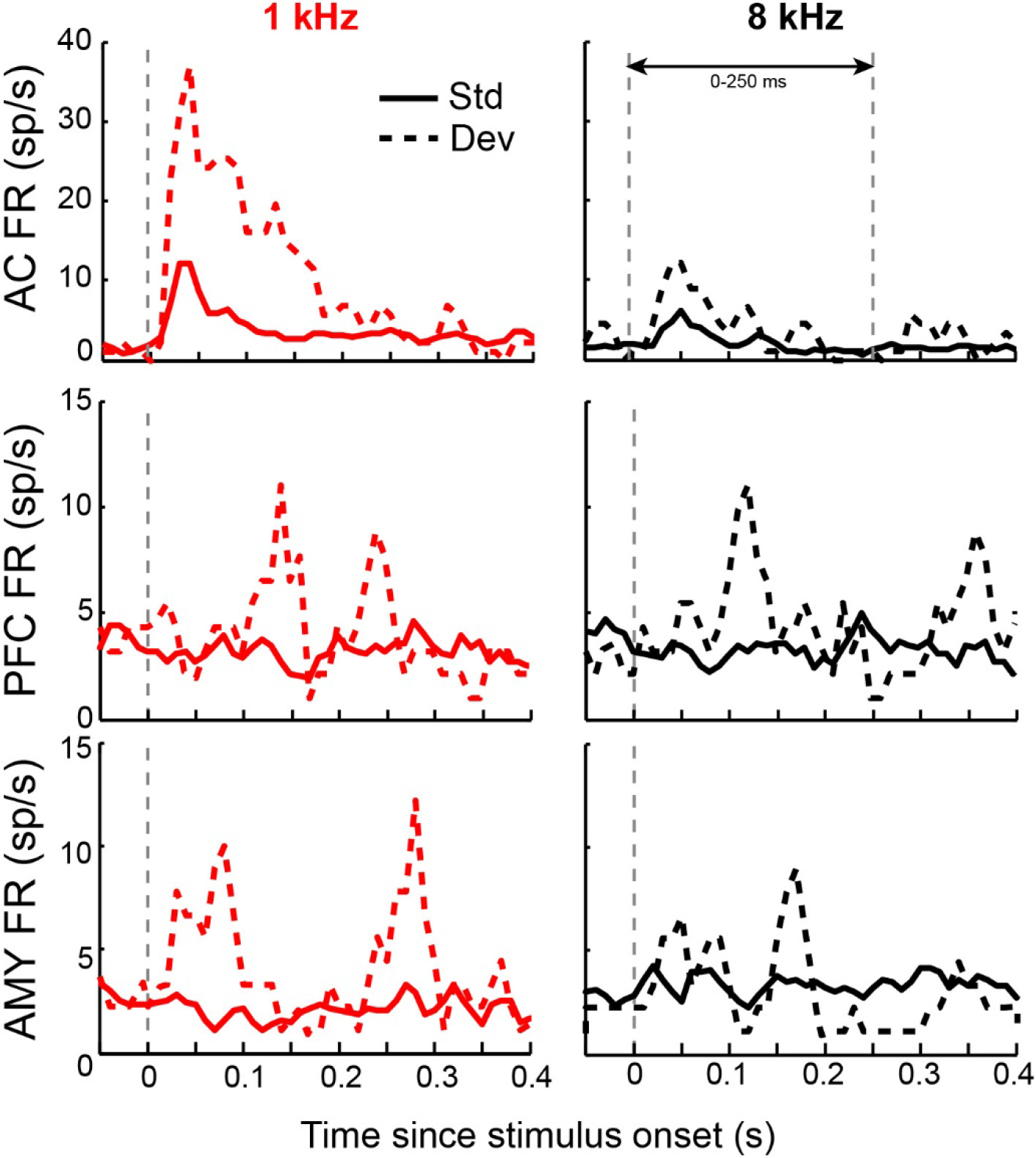
Example neuron responses. Peristimulus time histograms of average firing rate (FR) of an example single neuron response from each area (auditory cortex: AC, dorsolateral prefrontal cortex: PFC and lateral amygdala: AMY) to each stimulus and type combination. Each neuron shown is novelty selective. Note that in the prefrontal cortex and amygdala baseline firing rates and evoked activity are generally lower than in auditory cortex. PSTHs are smoothed for visualization purposes only.

The MMN measured at the scalp surface presumably represents activity of a neural population. While there is no direct mapping of neural activity to ERP components, a straightforward approach is to examine the magnitude of population activity of task responsive neurons and whether that population activity carries a novelty signal whether ththe novelty signal magnitude is different across areas). The magnitude of the response to the stimuli was approximately twice as strong in auditory cortex than in prefrontal cortex and amygdala (significant effect of area F(2,676)=7.2; p<0.001, between area post-hoc tests AC vs PFC: F(1,572)=7.4; p=0.006, AC vs AMY: F(2,676)=7.0; p=0.009, PFC vs AMY area: n.s., p=0.69). Auditory cortex responded more strongly to 1 kHz than to 8 kHz. Additionally, the magnitude of novelty signal, which can be measured as DEV-STD was dependent on the stimulus in auditory cortex and stronger for the 1 kHz than for the 8 kHz stimulus (Fig. 4). This stimulus selectivity was not present in prefrontal cortex and amygdala - novelty signals were the same magnitude irrespective of driving stimulus (Overall effect of type F(1,676)=58.2; p<0.001; AC stimulus F(1,468)=66.1, p<0.001; type F(1,468)=190.8, p<0.001; and stim x type F(1,468)=51.3, p<0.001; PFC type F(1,104)=41.0, p<0.001; AMY type F(1,461)=25.5, p<0.001 all other factors in PFC and amygdala p>0.1). Thus at a population level the average novelty signal in auditory cortex was robust and stimulus dependent, and in amygdala and prefrontal cortex it was smaller (though significant) and not stimulus dependent.

**Figure 3.**
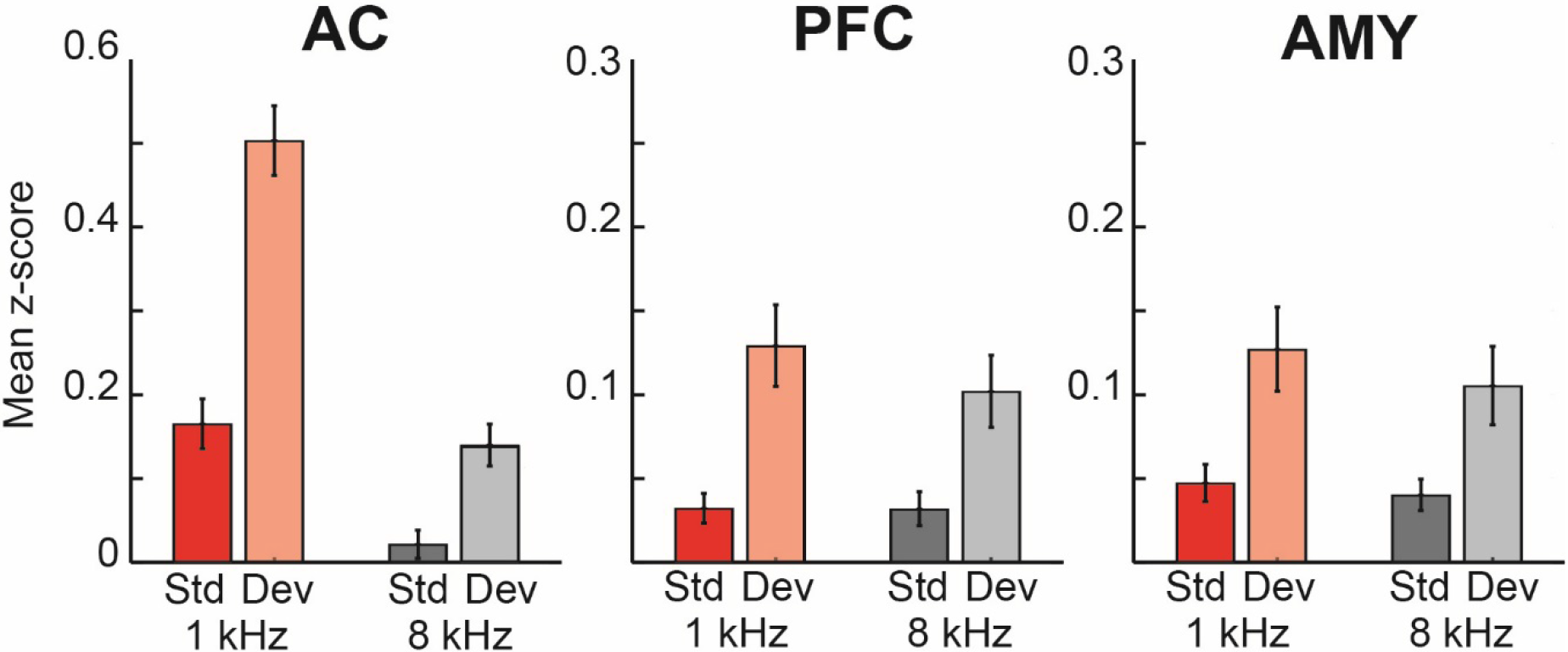
Population response by stimulus across areas. Average response magnitude for each stimulus, type (standard vs deviant), and area (AC n=462, PFC n=105, AMY n=105). Consistent with the exemplar responses in Fig 2, responses are on average larger in AC than PFC and AMY. Error bars are SEM across cells.

**Figure 4.**
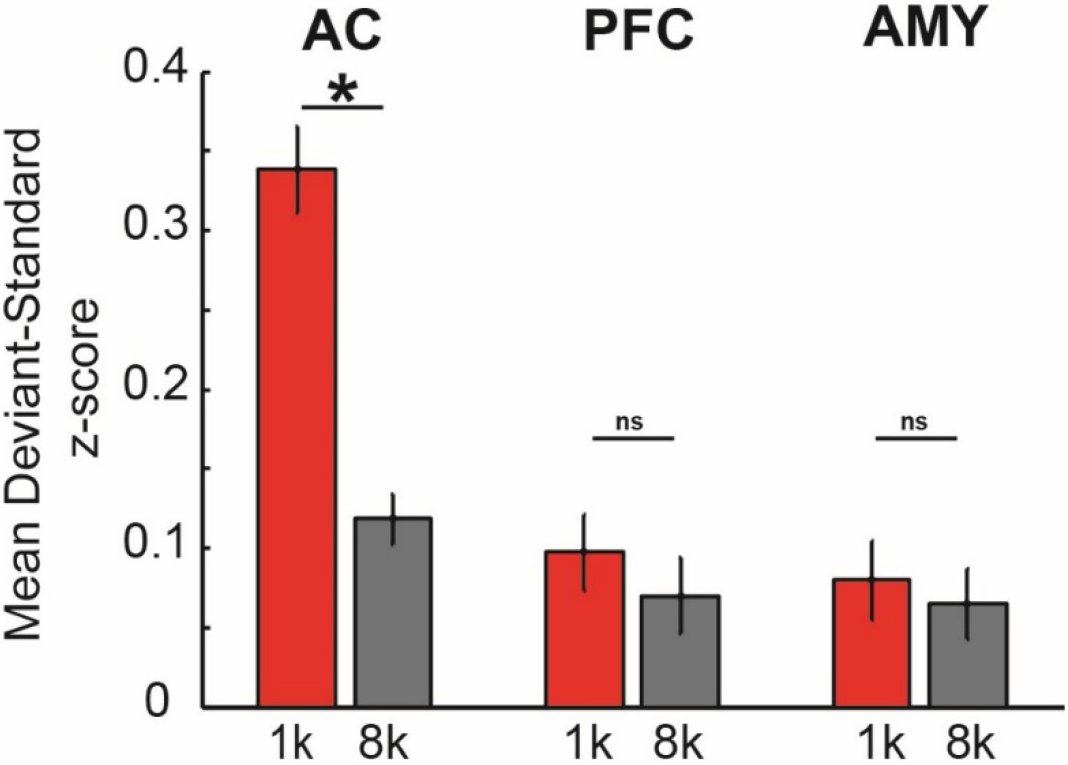
Population novelty signal across areas. Average novelty signal, computed as a neuron’s response to a stimulus as a standard subtracted from its response as a deviant (DEV-STD). Average magnitude of novelty signal is shown separately for each stimulus and area, error bars are SEM. All areas show a population-based novelty signal, but only auditory cortex shows a signal whose magnitude is affected by driving stimulus (sig difference denoted as a star).

For a more complete picture of what is driving this this population level signal, we next asked whether the novelty signal is driven by a few or many neurons in each area (e.g. a weak signal in prefrontal cortex could be a few neurons carrying a signal comparable in magnitude to that of auditory cortex or many neurons carrying a weak signal). In auditory cortex a substantial number of neurons show a novelty signal for at least one of the stimuli (166/462 = 35.9%). However, few single neurons in prefrontal cortex and amygdala showed a similar profile (PFC: 9/105 = 8.6%, AMY: 9/105 = 8.6%), and the magnitude of the signal is again smaller relative to that in auditory cortex. Thus, at the level of single neurons few neurons are carrying a strong signal in in the amygdala and prefrontal cortex - it is a weak signal carried by many that leads to the overall novelty population response. Note that again, these results were not dependent on window choice - they did not change substantially with smaller (0-100 ms) or larger (0-400 ms) fixed or sliding windows.

**Figure 5.**
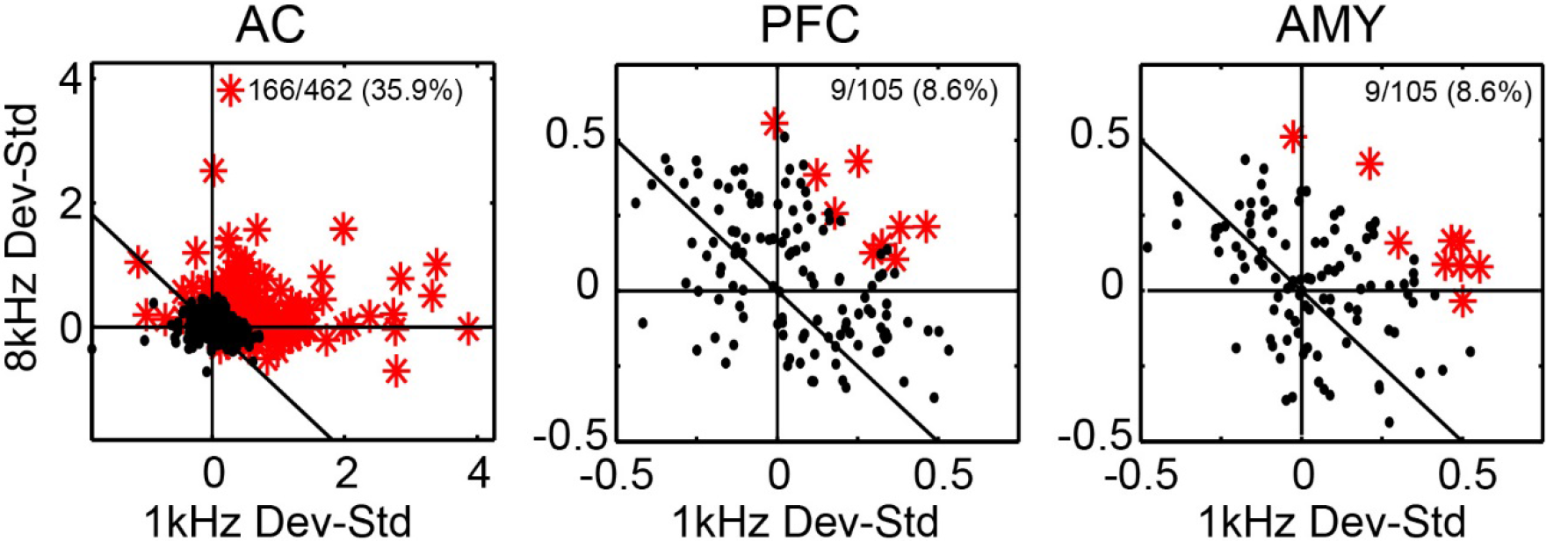
Individual neuron-based magnitude of novelty signal (response as deviant - response as standard) and novelty selectivity for each stimulus and area. Single neurons which exhibited a significant novelty signal are shown in red. This fraction is lower in PFC and AMY but greater than chance, and magnitude in these areas is lower than in AC (see text). Note that generally responses in AC are skewed towards stronger novelty responses for 1 kHz, but that there is less stimulus selectivity in PFC and AMY.

The next question is to compare the timecourse of the novelty signal between auditory cortex, prefrontal cortex and amygdala to see whether the novelty signal emerges first from sensory cortex. To do this, again we used two approaches, one based on population level activity and the second based at the level of single neurons, to more fully understand what is driving the population activity. As before, we first examine population based timecourses as an estimate of how novelty signal evolves in each area. We calculated the population average PSTHs for each stimulus x type combination over all the responsive neurons and examined when response to deviants differed from response to standards at a population level (Fig. 6A-C). In auditory cortex, the earliest difference was detected in the bin spanning 10-40 ms. In amygdala it was later, 30-60 ms. In prefrontal cortex, no 30 ms bin reached significance. The lack of an onset time in this analysis in prefrontal cortex probably due to the diffuse nature of the deviance signal in prefrontal cortex and was not due to the size of the analysis bin. We repeated the analysis with shorter and longer bins and prefrontal cortex did not reach significance in any bin from 10-150 ms. While this population approach is informative about when the signal generally is arising, it may obscure heterogeneous but important dynamics (e.g. a smaller population of prefrontal neurons respond before auditory cortex but the majority are later). To examine whether there are any fast dynamics that are obscured by averages, we analyzed latencies on a single-neuron basis. Here, we calculated the latency at which the deviance signal (e.g. selectivity to type) arose in each area on an individual neuron basis, using 30 ms bins to increase temporal resolution (Fig. 6A, as cumulative distribution functions). The 30 ms bins increases the overall number of neurons that carry a novelty signal in each area (relative to the fixed bins in Fig. 5 above), but importantly, as stated above, the overall trends between areas are the same. According to this single neuron analysis, sensitivity to type was still earlier in auditory cortex than amygdala and prefrontal cortex, and amygdala and PFC did not differ (AC=56.6 ms, n=332; PFC = 128.9 ms, n=36; AMY=137.2 ms, n=46; ANOVA sig for area F(2,411)=57.4; p<0.001, post hoc t-tests AC< PFC (t(366) = −7.6, p<0.001), AC<AMY t(376) = −9.1, p<0.001, PFC=AMY t(80) = 0.45, p = 0.64). Earliest novelty sensitivity latencies in auditory cortex were still well earlier than those in prefrontal cortex and amygdala. Taken together, these analyses suggest that a deviance signal occurs first in auditory cortex, and later emerges in prefrontal cortex and amygdala, and that the population signal is a veridical reflection of dynamics occurring at the single neuron level.

**Figure 6.**
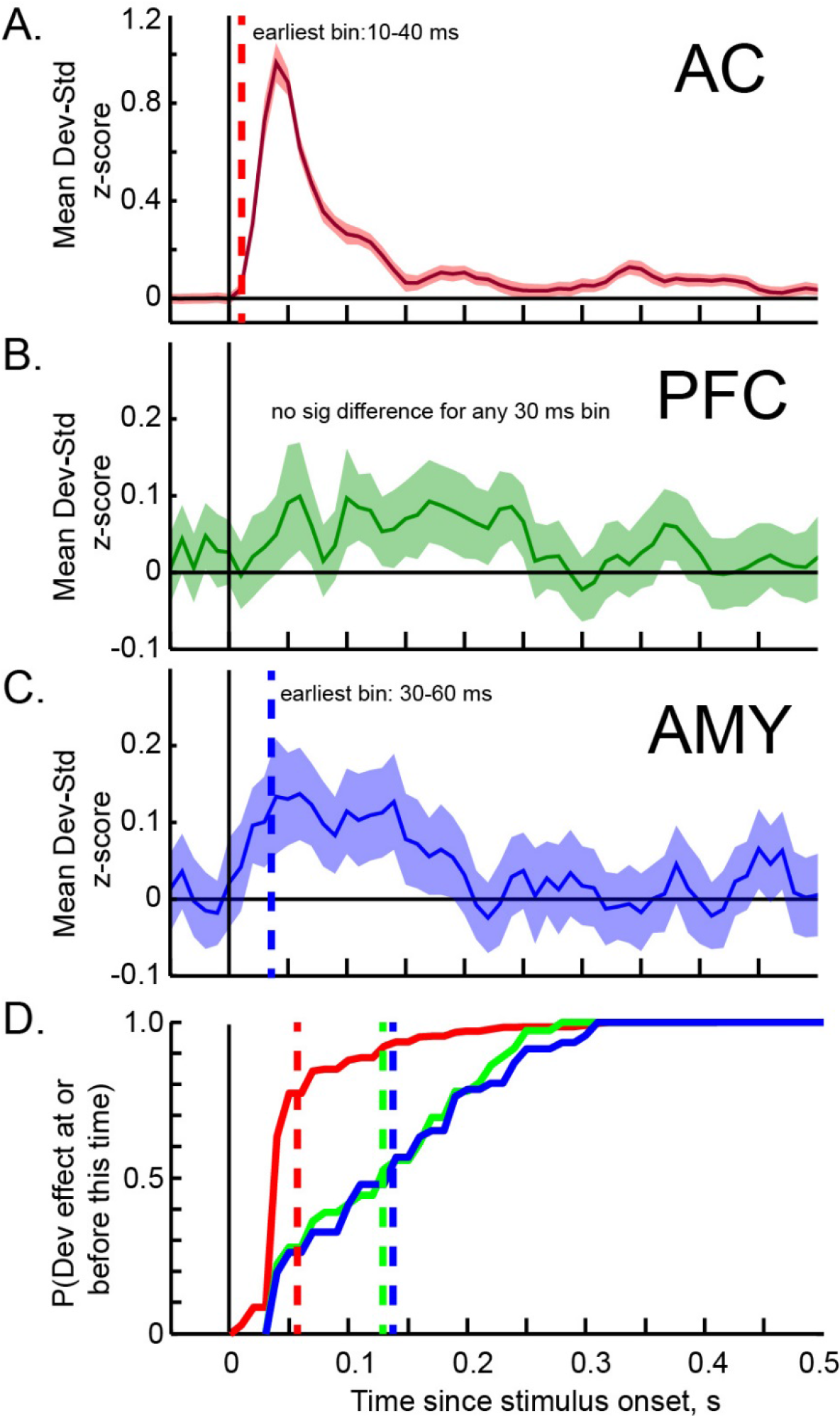
Latency of deviance signal across areas. A-C. Average PSTHs of DEV-STD differences for each area (shaded bars are SEM). Earliest 30 ms bin in which differences were significant denoted as vertical dotted line. Plots are smoothed for visualization purposes only. D. Cumulative distribution function of latencies of individual neuron sensitivity to stimulus type for each area. Mean of each area denoted as vertical dotted line.

As a final analysis to examine how the novelty signal evolves across areas of the brain, we examined the dynamics of response adaption across all three areas. Neural responses may adapt at the beginning of the block, and also after a deviant is presented, and the existence and time course of adaptation may change with level of processing and brain region. For example, responses that are strongly sensory driven may show strong adaptation effects, but one would not expect adaptation in higher-order responses that presumably reflect more generalized novelty processing. To address this, we analyzed adaption of the responses to standards and deviants in two ways. First we examined standard and deviant adaptation across blocks (Fig. 7A). Consistent within-block adaptation occurred in auditory cortex for three of the stimuli (mean and t-test on fit beta values: 1 kHz standard mean=−0.036, t(461)=− 10.9, p<0.001; 8 kHz standard mean=−0.016, t(461)=−4.7, p<0.001, 1 kHz deviant mean=−0.03, t(461)=− 4.2, p<0.016, 8 kHz deviant mean=−0.006, t-test ns, p=0.056). In auditory cortex, adaptation was faster for the standards and faster for the 1 kHz stimulus (2×2 stim x type within-neuron ANOVA stim F(1,900.2)=16.4, p<0.001, type F(1,472.5)=17.2, p<0.001, stim x type ns (p>0.1)). Adaptation was not observed in prefrontal cortex, and it was only observed in amygdala for the 8 kHz deviant (amygdala 8 kHz deviant mean=−0.02, t(104)=−3.6 p<0.001, all other p>0.1). Note that this within block adaptation did not contribute to the novelty signal described above, as the first deviant was not presented until at least 10 standards had been presented (dotted line), after the within-block adaptation had occurred. A second type of adaptation is the effect that the deviant has on subsequent standards (Fig 7B). We found that post-deviant adaptation occurred in auditory cortex for the 1 kHz stimulus (mean=−0.02, t(461)=−3.3, p<0.001), but not for other stimuli or areas (all other t-tests p>0.1). Note that the lower responses in amygdala and prefrontal cortex, as well as auditory cortex 8 kHz, may make adaptation difficult to detect. Generally however, it appears that stimulus-specific characteristics of adaptation seen in auditory cortex were not present in prefrontal cortex or amygdala, consistent with the more generalized novelty signal these higher-order areas appear to carry.

**Figure 7.**
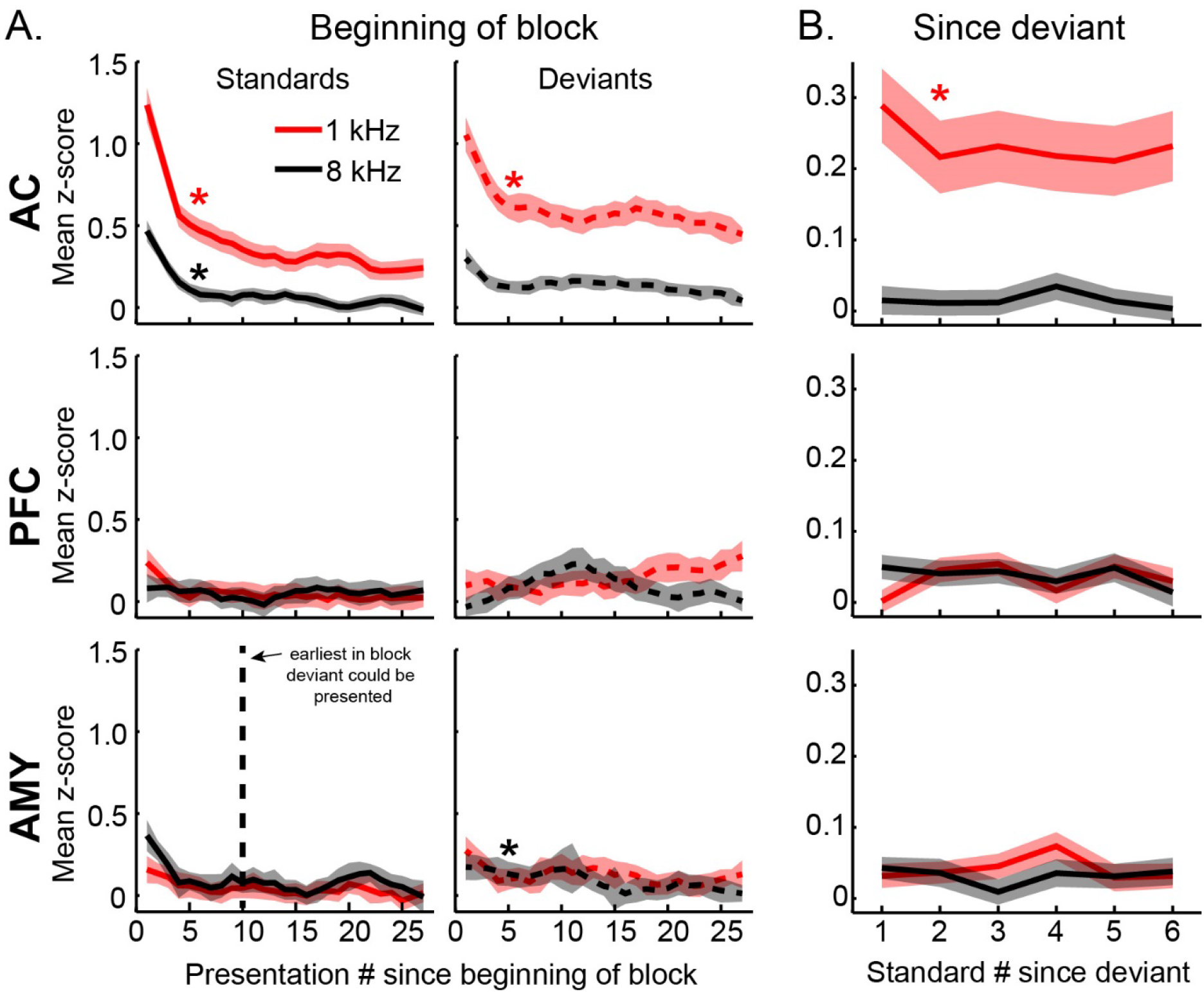
Adaptation effects across areas. A. Adaptation since block start. Average response to a standard (solid) and deviant (dotted) plotted by ordinal presentation number since beginning of block for each area, stimulus, and type (shaded bars are SEM). Significant adaptation is denoted with a star. Vertical dotted line indicates the earliest that a deviant could be presented within a block (all blocks started with >10 standards). B. Adaptation since deviant. Average response to standard after presentation of a deviant for each area and stimulus (shaded bars are SEM). Plots are smoothed for visualization purposes only.

## Discussion

The overall goal of this experiment was to leverage the spatial and temporal specificity of a primate model to address outstanding questions about novelty processing. Specifically, we compared the magnitude, timing, and stimulus specificity of auditory oddball-based novelty signals in auditory cortex, dorsolateral prefrontal cortex, and the basolateral amygdala. These data were recorded under identical conditions from a substantial number of neurons in the alert macaque, so effects could be captured and compared. This approach allows us to test predictions made by the predictive error/sensory memory model of the MMN, which posits that a frontal generator exists which is sensitive to a change in the expected sensory stream, and that this signal should be both abstracted from and later than the one in auditory cortex (Garrido et al., 2009; Giard et al., 1990; Naatanen et al., 1992). We find that, compared to the novelty signal seen in auditory cortex, the signal seen in dorsolateral prefrontal cortex and basolateral amygdala is smaller in magnitude, longer in latency, and not stimulus dependent. This is generally consistent with predictions of the predictive error hypothesis in that novelty signals in prefrontal cortex should be generally later than in auditory cortex, as well as abstracted from stimulus specific effects seen in auditory cortex. However, the fact that signals in amygdala were comparable in magnitude and timing to those in prefrontal cortex, and both prefrontal and amygdala signals were generally much weaker than those in auditory cortex, is not wholly consistent with the account.

The approach we present here allows more precise cross area comparisons than previous noninvasive or ECoG approaches. We see that novelty response in prefrontal cortex and amygdala has properties that are strikingly similar to each other yet do not appear to be simply inherited from the signal seen in auditory cortex. First, the stimulus specificity seen in auditory cortex was no longer significant in prefrontal cortex or the amygdala. Second, the within-block adaptation and post-deviance adaptation seen in auditory cortex were also no longer observed. This suggests that prefrontal cortex and amygdala have, to some degree, an abstracted deviance signal distinct from auditory cortex. These areas signal “difference”, but they do not reflect low level properties of the driving stimuli (e.g. spectral overlap). The timing of the difference signal is also consistent with that interpretation. The prefrontal difference signal lags that of auditory cortex by an average of ~100 ms, and the amygdala difference signal lags auditory cortex by between 20-100 ms. These observations place useful quantitative constraints on putative generators of the auditory oddball-based MMN, and indicate that subcortical areas, such as the amygdala, may need to be included in future explanatory accounts of auditory oddball.

Previously there have been only indirect measures suggesting that frontal MMN generators are later than sensory generators (Deouell, 2007; Rinne et al., 2000; Tse & Penney, 2008). Human intracranial studies of auditory oddball paradigms have supported the existence of separate frontal and temporal components (Durschmid, Edwards, et al., 2016; Edwards et al., 2005; El Karoui et al., 2015; Kropotov et al., 2000; Liasis, Towell, Alho, & Boyd, 2001; Rosburg et al., 2005). However, due to the heterogeneity of the clinical placement of monitoring electrodes within and between patients, it has been difficult to compare the magnitude and timing of the deviance signal between these areas. In addition, our study extends our understanding of putative substrates of the oddball-based novelty processing to the amygdala. This builds on a hypothesized role of the amygdala in salience detection (Blackford et al., 2010). Here, we show that amygdala correlates of novelty are comparable in magnitude, timing, and response characteristics to those seen in prefrontal cortex. Previous accounts may have been biased towards cortical generators because noninvasive scalp-based measures are less sensitive to deep generators. This new finding may require an update of the accounts of the generators underlying the MMN.

Experiments in both rodents (Parras et al., 2017; Taaseh, Yaron, & Nelken, 2011; Ulanovsky et al., 2003; Yarden & Nelken, 2017) and primates (Fishman, 2014; Fishman & Steinschneider, 2012; Javitt et al., 1994) have examined single neuron correlates of the MMN in the early auditory hierarchy (cochlear nucleus through auditory cortex). The novelty signal in auditory cortex that we observed in this study had a magnitude, time course, and stimulus dependent characteristics consistent with what has been seen in population-based analyses of single neuron studies in auditory cortex (Farley, Quirk, Doherty, & Christian, 2010; Nieto-Diego & Malmierca, 2016; Parras et al., 2017; Ulanovsky et al., 2004; Ulanovsky et al., 2003). There are two methodological differences worth noting between our study and these studies, driven by the aim of our study, which was to compare the characteristics and timing of two putative higher-order generators of the deviance signal, to the deviance signal seen in auditory cortex. First, the stimuli used in this study were wideband and kept constant for all of the data collected, instead of the previous studies’ method of using pure tones whose identity is chosen to carefully flank the individual tuning of the neurons studied (and “population” responses collapsed across stimuli later). Thus we would expect the population responses in our study to be more comparable to human studies in which all the data are collected using the same stimuli for all of the areas. Second, the majority of the single neuron studies were performed under anesthesia (but see Farley et al., 2010; Parras et al., 2017). While deviance detection, especially the mismatch negativity, can be elicited under anesthesia, the anesthetic state may affect higher order areas and minimize top-down effects seen from, for example, prefrontal cortex, which does not exhibit robust responses under anesthesia.

Utilizing the spatial and temporal specificity of a macaque model, this approach overcomes limitations in our understanding from earlier findings derived from noninvasive methods and human intraoperative recordings. There are multiple aspects of these findings that merit further investigation. The MMN-indexed deviance signal is thought to contain an adaptation component (suppression of the expected), and/or a surprise component (enhancement of the unexpected). Note that detection of deviance is of great evolutionary importance and therefore it is likely supported by a broad network of areas whose magnitude, recruitment, and mechanism (e.g. adaptation vs surprise) may depend on task type and behavioral requirements (MacLean, Blundon, & Ward, 2015; Warbrick, Reske, & Shah, 2013). Nevertheless, even in the “simple” auditory oddball paradigm, the relative contributions of each of these mechanisms (adaptation vs surprise) to the recorded signal has been a matter of intense debate, especially in the early auditory hierarchy (Fishman, 2014; Parras et al., 2017). While the “error prediction” model posits that the frontal activation is an important generator of the MMN, the “adaptation only” hypothesis (.e.g Fishman, 2014), would posit that it is epiphenomenal. Our study finds that the signals in amygdala and prefrontal cortex have characteristics described by the error prediction hypothesis, but that the strength in both areas is weaker than that of auditory cortex. Therefore, an important future direction will be to establish causal links between prefrontal cortex and amygdala and the deviance signal seen in auditory cortex and at the scalp to be able to disambiguate between these accounts. An earlier study reports that patients with prefrontal lesions exhibit a smaller scalp-measured mismatch negativity (Alain, Woods, & Knight, 1998; Alho, Woods, Algazi, Knight, & Naatanen, 1994). Though this would seem to imply a causal relationship of the prefrontal cortex to the scalp-recorded MMN, it is difficult to compare the magnitude of ERP components when there is substantial difference in damage to the underlying cortex. Due to its shared primate homology and established MMN (Gil-da-Costa et al., 2013) the macaque model is an ideal model to test these questions using reversible inactivation. Towards that end, these data provide a significant advance in our conceptual framework of how deviance is processed outside of the early subcortical to cortical auditory hierarchy and open new directions of investigation in one of the dominant deviance detection paradigms, the auditory oddball paradigm.

## Acknowledgments

The authors would like to thank Dr. Israel Nelken and Dr. Brian Scott for valuable feedback, Dr. Richard Saunders for surgical assistance, Anna Leigh Brown and Jess Jacobs for assistance in data collection, and the NIH Section on Instrumentation assisted in custom manufacture of recording chambers and grid Research was supported by NIMH DIRP ZIA MH002928-01 to BA and ZIA MH001101-25 to MM.

